# Sound lateralization Ability is affected by saccade direction but not Eye Movement-Related Eardrum Oscillations (EMREOs)

**DOI:** 10.1101/2025.11.05.686724

**Authors:** Nancy Sotero Silva, Felix Bröhl, Christoph Kayser

## Abstract

Eye-movement-related eardrum oscillations (EMREOs) are pressure changes recorded in the ear that supposedly reflect displacements of the tympanic membrane induced by saccadic eye movements. Previous studies hypothesized that the underlying mechanisms might play a role in combining visual and acoustic spatial information. Yet, whether and how the eardrum moves during an EMREO and whether this movement affects acoustic spatial perception remains unclear. We here probed human acoustic lateralization performance for sounds presented at different times during a saccade (hence the EMREO) in two tasks, one relying on free-field sounds and one presenting sounds in-ear. Since the EMREO generation likely involves the middle ear muscles, whose tension can alter sound transmission, it is possible that judgements of sound locations may vary with the state of the ERMEO at the time of sound presentation. However, when testing two specific hypotheses of how movements of the eardrum underlying the EMREO may affect spatial hearing, we found no evidence in support of this. Still, and in line with previous studies, we found that participants’ lateralization responses were shaped by the spatial congruency of the saccade target direction and the sound direction. Thus, either the eardrum does not move directly as reflected by the EMREO signal, or despite its movement the underlying changes at the tympanic membrane only have minimal perceptual impact. Our results call for more refined studies to understand how the eardrum moves during a saccade and whether or how the EMREO impacts spatial perception.

## Introduction

Many objects in real life are multisensory, and our brain needs to combine the stimuli gathered from different sensory modalities to estimate their location. This, for example, requires the combination of spatial signals received by the eyes and ears. Recent work highlights the possibility that this process may even engage the periphery of the auditory pathways: studies have described so-called eye movement-related eardrum oscillations (EMREOs), low-frequency signals recorded using in-ear microphones that are precisely aligned to saccade onsets (Sotero Silva et al., 2025; King et al., 2023; Lovich et al., 2023a; Gruters et al., 2018). These EMREO signals last for about 50 to 100 ms and comprise multiple deflections whose sign relates to the direction of the inducing saccade (Murphy et al., 2020; Lovich et al., 2023a, 2023b, King et al., 2023; Bröhl and Kayser, 2023).

The EMREOs elicited by saccades to the same or opposite directions to the ear under study exhibit deflections of opposing sign but also feature distinct time courses: the EMREOs in the two ears tend to move in opposition but not in strict anti-phase. Starting from the assumption that the EMREO is indeed indicative of the actual movement of the tympanic membrane, this suggests that the left and right eardrums move differentially during a given saccade (Sotero Silva et al., 2025; Bröhl and Kayser, 2023; Gruters et al., 20218). Hence, sounds presented at different times relative to a saccade are received with the two tympanic membranes deflected in- and outwards and with different amplitudes. This may differentially affect the transduction of acoustic cues relevant for the interpretation of a sound source’s location by the auditory system, a process that relies on the coordinated information from the two ears (Culling and Akeroyd, 2010). Support for such an impact of EMREOs on sound transduction comes from a study showing that the contraction of the middle ear muscles changes the shape of the tympanic membrane and alters low-frequency sound transmission delays (i.e., the time for the response of the stapes after arrival of a sound at the ear canal) (Cho, Ravicz and Puria, 2023). Thus, it is possible that saccades affect the transduction of acoustic cues in a manner dependent on the EMREO time course, for example by affecting the encoding of acoustic features relevant for precise auditory lateralization (to resolve the direction - right or left - of the sound source relative to the center of the head), such as interaural time (ITD) and level differences (ILD) (Cho, Ravicz and Puria, 2023; Lovich et al., 2023a, Tasko et al., 2022). As a result, sounds received during larger EMREO deflections may be transduced less faithfully than sounds during smaller deflections, resulting in an influence of the magnitude of the EMREO signal on the accuracy of spatial estimates. In addition, or alternatively, sounds received during EMREO deflections of opposing signs may be transduced differently, resulting in an influence of the sign of the EMREO on systematic directional biases in auditory spatial judgements.

However, the actual influence of eardrum movements as reflected by the EMREO signal on auditory perception remains little investigated. In a previous study we have examined this in a near-threshold sound detection task: we found that the time when a sound was presented relative to the EMREO did not have a significant effect on detection performance (Bröhl and Kayser, 2023). Nevertheless, this negative result may simply reflect the specific nature of the task, which did not require participants to exploit the spatial information carried by a sound, but simply asked them to detect whether a sound was heard over the alternative that no sound was heard. Accordingly, the experiment did not test whether the quality of the transduced sound energy was altered by its relative timing to the EMREO. Still, such an impact of EMREOs on auditory spatial judgments has not been tested so far. We hence asked whether the timing of a sound relative to the EMREO affects how its location can be judged. For this we tested human participants in two tasks requiring them to resolve the spatial information about a sound in order to judge its source. These tasks required the comparison of the relative horizontal location of two subsequent tones that were either presented by loudspeakers (free-field task) or were delivered using in-ear phones at varying inter-aural time differences (in-ear task). Based on the spatial information derived from each sound, participants had to decide whether the second tone was lateralized to the left or right relative to the first tone.

Because EMREOs are triggered by saccadic eye movements, any investigation of how they influence auditory spatial perception must also address the relation between spatial hearing and oculomotor behavior in general. Previous studies have reported that saccades do not impair basic auditory abilities such as sound detection or pitch discrimination (Harris and Lieberman,□1996; Paire et al., 2025). However, studies using spatial hearing tasks (i.e., addressing acoustic localization and/or lateralization) have demonstrated that sounds are misjudged during static fixations: when participants fixate an eccentric visual target their estimates of horizontal sound locations are biased, though the direction of bias seems to differ between experimental paradigms, which vary considerably in terms of how responses to acoustic tasks are recorded (Lewald and Ehrenstein, 1996; Lewald, 1997; Getzmann, 2002; Klingenhoefer and Bremmer, 2009; Van Grootel and Van Opstal, 2010; Razavi et al., 2007). Saccades, in contrast, seem to have a smaller effect on spatial hearing: the estimates of sound locations just prior to or during a saccade are not affected differentially by the time course of the saccade (Klingenhoefer and Bremmer, 2009). Overall, perceived sound locations seem to be biased towards visual saccade targets independently of the timing of the relevant tones relative to saccade onset (Klingelhoefer and Bremmer, 2009; Pavani et al., 2008; Binda et al., 2007). Similarly, estimates of auditory motion were reported to be biased in the direction of a saccade (Krüger et al., 2016), suggesting that the relative congruency of saccade direction and relative sound locations may shape how a sound’s location is judged. Translated to the present experimental paradigm, in which participants compared the relative horizontal location of two subsequent sounds, we included the congruency of saccade direction and horizontal sound locations (thus, its direction) as a factor in the analysis. These results predict that participants may judge the relative sound locations in accordance with the saccade direction.

To probe an influence of EMREOs on spatial hearing, the present study tested the same participants performing two auditory spatial discrimination tasks for sounds presented just prior to or during a visually-guided horizontal saccade. Each trial required the comparison of the perceived horizontal location of two subsequent sounds: the first sound was always presented centrally and served as a reference, while the second sound was located horizontally to the left or right, and participants had to judge this sounds’ perceived direction as either left or right to the first one. Importantly, we jittered the presentation times of the sounds relative to saccade onset, so that the presentation of the second sound randomly fell into different phases of the concurrently recorded EMREOs. To test whether participants’ spatial hearing was affected by the timing of the sound relative to the EMREO, we analyzed the data for two specific questions: 1) whether the direction of the EMREO deflection in each ear induces a systematic bias in lateralization judgments, and 2) whether the magnitude of the EMREO signal (and presumably of the eardrum displacement) affects how well a sound location can be judged. Overall, our results speak against any such effect in either of the two experiments.

## Materials and methods

### Participants

The data presented here were collected as part of a two-experiment study. The first part of that study was designed to characterize EMREOs during visually- and acoustically-guided saccades and has been published (Sotero Silva et al., 2025). The second part is reported here. 34 participants were recruited among the university and community members (24 females, 10 males, mean ± SD age = 27,3 ± 5,3 years) without self-reported hearing or vision impairments and neurological or psychiatric disorders. Each participant provided written informed consent and received 15 euros per hour as compensation. The study was approved by the ethics committee of Bielefeld University (#2021-218). Initially, participants were visually screened for obstructions in their ear canals and performed pure tone audiometry to establish air conduction hearing thresholds. They underwent tympanometry and were tested for acoustic reflexes at 500, 1000, 2000 and 4000 Hz. Individuals with an audiometry quadritone average threshold below 20 dB were considered to have normal hearing (World Health Organization, 2020) and were included in the actual experiment. Data was obtained in two sessions taking place on two different days, with each session testing one of the two experiments described below. We ensured that the functioning of the ear was comparable between sessions based on the hearing screening and middle ear assessments.

### Experimental Setup

The experiments were conducted in the same set-up as described in Sotero Silva et al. (2025). Recordings took place in a soundproofed and electrically shielded booth with no light sources other than a projector. Participants remained seated in front of a table with their head on a chinrest, aligned to the center of a sound-permeable screen 100 cm away (Screen International Modigliani, 2 × 1 m). In-ear microphone recordings were collected using two Etymotic ER-10C (Etymotic Research) systems, via probes that served both for sound presentation and microphone recording (we name the phones/microphone assembly as “probes” when we refer to the device in general; as “in-ear phones” to indicate the stimuli presentation; and as “microphone” to indicate the recordings). Microphone signals were pre-amplified using the ER-10C DPOAE amplifiers (Etymotic Research), with gain set to +40 dB. The signal was digitized through an ActiveTwo AD-box (BioSemi) at a sampling rate of 2048 Hz. Visual stimuli were projected (Acer Predator Z650, Acer; 60 Hz refresh rate) onto the screen. The acoustic stimuli were created using Sound Blaster Soundcards with sampling rate 44100 Hz and were presented differently depending on the experiment. In one experiment they came from an array of 5 free-field speakers (Monacor MKS-26/SW, MONACOR) located behind the screen and positioned between -22 and 22 degrees (spaced in 11 degrees steps) azimuth and 8 cm from the screen border (around 108 cm away from the participant’s head). In the other experiment they were amplified with a HeadAMP 4 (ARTcessories) amplifier and presented via the ER-10C in-ear phones. The Psychophysics toolbox (Brainard, 1997) for MATLAB (The MathWorks, 2022) was used to control the stimulus presentation, which was synchronized to the recording system using TTL pulses. Figure 1A represents the experimental set-up.

**Figure 1.**
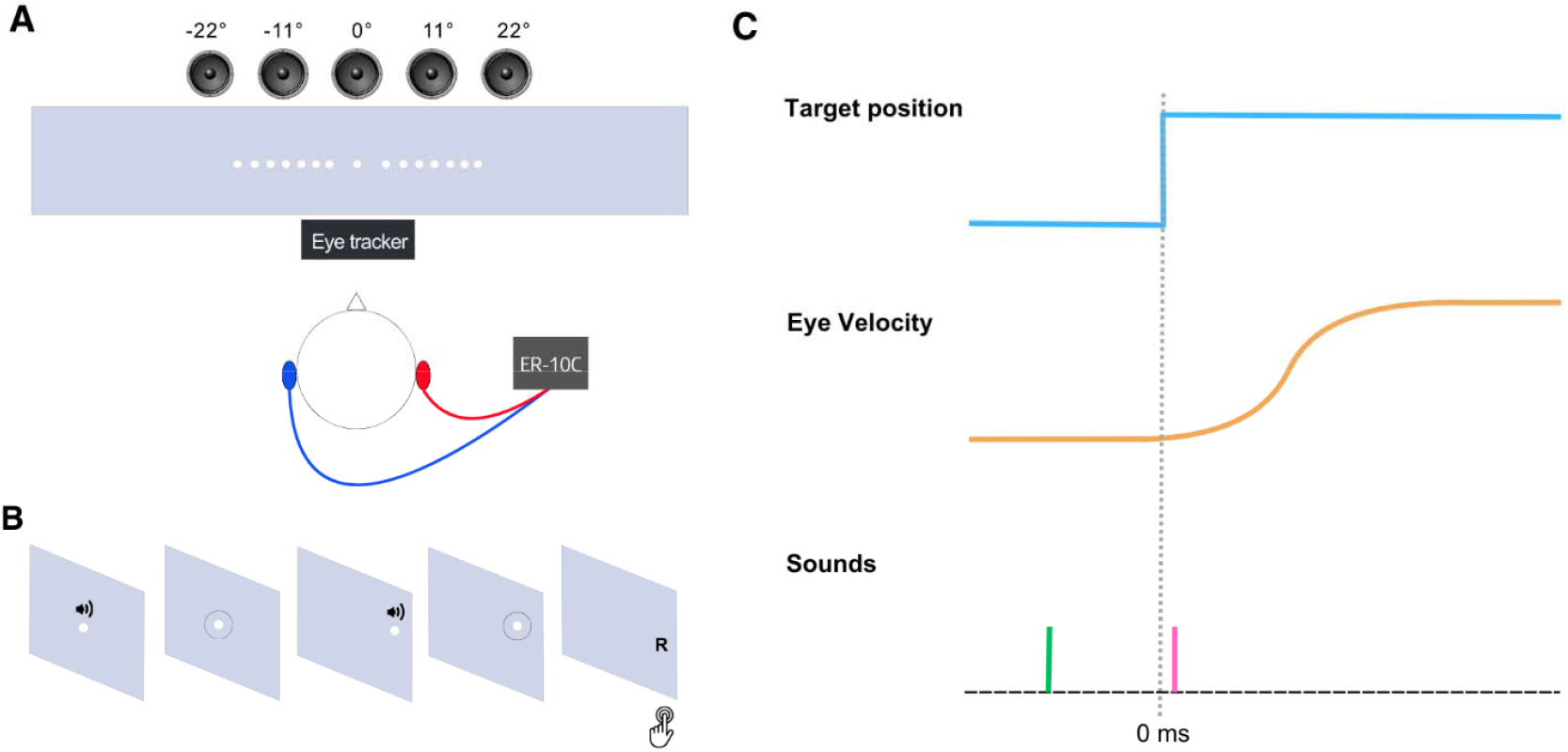
Experimental Paradigm and Design. (**A**) shows a representation of the experimental setup, indicating the position of the participant, the screen for the projection of the visual targets, the eye tracker and the speakers positioned behind the screen. In-ear microphones for the left (blue) and right (red) connected to the ER-10C amplifier are also displayed. (**B**) shows the sequence of events. The white dot indicates the fixation target, the speaker icon the positions of the two tones. The letter “R” represents the expected response for the sound lateralization task (in this example, “right”, obtained by arrow keys press). (**C**) Schematic timeline for one trial. The visual saccade target (light blue) jumps from 0 (fixation period) to the target location, the eye follows subsequently (eye velocity; orange). Tones were presented around the fixation (first tone, in green) and during the saccade (second tone, in pink). The dotted gray line represents the saccade onset (0 ms).

Eye movements were recorded at 1000 Hz from the left eye using a desktop mounted infrared-based EyeLink 1000 Plus Eye Tracker (SR Research Ltd., 2015) running Host Software version 5.15. Eye-tracking calibration and validation was performed for each block using the built-in 5-point grid. The parameters for saccade detection in the EyeLink system (“cognitive” setting) were a velocity threshold of 30°/s and an acceleration threshold of 8000°/s. With this standard criterion for saccade detection, the actual saccade velocity already ramps a few milliseconds prior to the detected onset, as visible in the group-data of saccade velocity (Figure 3B). Eye positions were recorded as horizontal and vertical coordinates in degrees of visual angle.

To ensure the quality of the microphone and in-ear recordings control measurements were performed at the beginning of each session. First a recording was made in the empty booth. Microphone responses for 15 seconds of silence followed by 20 tone stimuli presented from the speakers (10 stimuli at 100 Hz, 10 stimuli at 200 Hz, duration 400 ms) were measured. Secondly, after the insertion of the probes in the participants’ ears, the same measurement was repeated. Adjustments for the microphone insertion and signal corrections across ears for data analysis were performed based on these measurements as described in Sotero Silva et al. (2025).

### Experimental Paradigms

During each session we first obtained data during visually- and auditory-guided saccades as described in our previous study (Sotero Silva et al., 2025). Subsequently, participants were instructed about the respective acoustic tasks, performed training trials, completed a threshold estimation procedure, and then performed four blocks of the actual tasks. Importantly, the acoustic tasks were designed around each participant’s individual perceptual threshold to avoid near ceiling performance that would make the detection of EMREO-related effects on spatial hearing difficult.

Each trial involved the presentation of two tones (1200 Hz 15 ms sine burst, cosine ramped and presented at 65 dB SPL; inter-sound interval of 240 ms +/-20 ms jitter) presented either from the free-field speakers (“free-field task”) or using varying ITDs presented via the in-ear phones (“in-ear task”; see Figure 1A). For the free-field task, a continuous sound location was achieved by linearly interpolating sound amplitudes between neighboring speakers. For the in-ear task, sounds were delivered via the ER-10C systems and were lateralized by adjusting their relative ITDs to achieve the required perceptual auditory location (sound locations were converted from azimuth angles into ITD values based on a generic head-radius function of 8.76 cm). The first tone was always presented in the center location, and served as a reference stimulus (“Tone 1”). The second tone was presented either to the left or right of the first one and was the probe (“Tone 2”). Participants’ task was a lateralization judgement by which they had to indicate whether the second tone was “perceived to the right or left relative to the first tone” by pressing one of two response buttons (Figure 1B). We refer to “lateralization” as the auditory processing ability related to the interpretation of the sound source’s spatial features’ for defining its location in terms of left or right (American Speech-Language-Hearing Association, 2005; Culling and Akeroyd, 2010). Thus, we use the term lateralization to refer to the subject’s performance in the task and refer to “sound direction” as the spatial difference in the two sounds (left/right) in the following text. During the familiarization phase, participants were initially presented with the tones together with an indication of the correct response on the screen. In this phase, the azimuthal distance between the locations of the first and the second tones remained constant (-6 degrees for the left, 6 degrees for the right). Subsequently they had to perform the task without feedback. Once they reached 70% correct responses, they performed a block to determine their Minimum Audible Angle (MAA). This was measured using three interleaved 3-down 1-up staircases from which we averaged the individual MAA’s to obtain the participants’ threshold (group-level values: mean = 3.99 deg, SD= 0.67 deg for the free-field task; mean = 3.48 deg, SD= 1.67 deg for the in-ear task). For the actual task we then presented tones with a location difference near threshold, adding a trial-wise random jitter to the participant-wise MAA for each trial (drawn from a uniform interval spanning +/-10% of the respective MAA), allowing some variety in the locations for Tone 2. Participants completed 4 blocks of 150 trials for each task.

During the actual tasks participants had to perform both visually-guided saccades and the acoustic discrimination task. Visually-guided saccades were cued using a white dot (0.2-degree radius). The dot first appeared in the center of the screen for 400 ms and participants had to fixate this. Subsequently, the fixation dot jumped to a random horizontal location to the right or the left (target locations were between -9 and -3 degrees to the left, and +3 and +9 degrees azimuth to the right, in one-degree steps) and participants were instructed to follow this dot as quickly as possible. The dot remained present until detection of a saccade or for 500 ms if a saccade was not detected during this time. Note that participants had already performed similar visually-guided saccades in a separate block designed to just analyze EMREO properties (Sotero Silva et al., 2025) and were well familiar with this task. Inter-trial intervals were randomly sampled from the interval between 900 to 1400 ms.

The timing of the two sounds was adjusted to the saccade behavior of each participant with the aim to sample different locations during the saccade-induced EMREO with the second tone (Figure 1C). To achieve this, we estimated the average saccade delay and its variability for each participant based on the separately obtained visual saccade task. Based on this we derived an estimate of when to present the sequence of tones, which also included estimates of the delays between auditory and visual presentation times in the setup. For most participants the average saccade delay was on the order of 250 ms and hence the first tone was usually presented prior to the saccade and the second tone during the saccade. However, given the trial-to-trial variability of saccade latencies, this resulted in some of the second tones to be presented prior to the EMREO, some during, and some after the EMREO. Each trial then comprised a horizontal visually-guided saccade starting from a central fixation dot and two tones. Example data for one participant are shown in Figure 3A, which illustrates the trial-wise timing of the saccade target, and the presentation times of Tones 1 and 2 respectively. The direction of the instructed saccade and the direction of the relative sound locations were pseudo-randomized. As a result, the relative location between Tone 1 and 2 could match (termed congruent) the direction of the saccade or be opposed to this (termed incongruent).

### Processing of EMREO data

For each task and participant, we computed the trial-averaged EMREOs separately for each ear and each saccade direction, as in previous work (Sotero Silva et al., 2025; Bröhl and Kayser, 2023). We aligned the microphone recordings to the onset of the largest saccade following the jump of the fixation dot. To avoid artifacts, we only included saccades whose amplitude was within the range 2 to 16 degrees visual angle, for which the initial fixation prior to the saccade was stable, and for which the entire horizontal and vertical traces of the eye position were contained in an 18 x 18 degrees window around the central fixation point. These EMREO epochs were z-scored and screened for extreme values. Epochs for which the maximal Z-score of the EMREO signal at any point in time exceeded 4.5 SDs of the overall signal were removed. We also visually inspected the participant-wise EMREOs and removed recordings presenting clear contamination with external noise or artifacts. From the data obtained from 34 participants, we retained datasets of 26 participants from the free-field task and of 21 participants from the in-ear task for further analysis based on the quality of the EMREO signal. The resulting trial-averaged EMREOs are shown for the group-data in Figure 3C and D and illustrate the opposing deflections for saccades grouped by direction in each ear.

### Analysis of behavioral data

We used regression models to test how the sound direction, the direction of the saccade and the relative timing of the tones to saccade onset (termed “latency” below) affected participants’ lateralization behavior. These models were fit to the trial-wise data to exploit the full variability offered by the large number of experimental trials collected. These models were used to predict three metrics of behavior: response times, response accuracy and the actual left/right responses (termed response “choice”). While the analysis of response accuracy focuses on how well participants discriminated between the two potential sound directions, the analysis of choice directly captures how sound direction and saccade direction shape participants’ lateralization behavior. We fit those models to the collective single-trial data from all participants, including participants as random effects. The models for response times were fit after transforming these (using a square-root transform) and were fit using assumptions of normal distributions and an identity link function. Models for accuracy and choice were fit using assumptions of binomial variables and logit link functions. As predictors, we included sound direction, saccade direction, and the latency of the tone. Since we expected that participants’ behavior may depend on the congruence of sound direction and saccade direction, we also included the interaction between these factors. For accuracy, the complete model would be as follows, with “Sound direction” being the direction of Tone 2 (left/right), Saccade Direction the direction of the saccade (left/right), and latency in milliseconds:

Accuracy ∼ Sound + Saccade + Sound:Saccade + Latency + (1|Participant).

We used these models in two ways: first, to estimate the predictive power of a given variable of interest and second to estimate the significance of the slope (regression beta) of individual predictors. The predictive power was estimated using model comparison. For this we contrasted a model including the predictor of interest (e.g. saccade direction) with a model excluding this predictor (Wagenmakers, 2007). For each model we extracted the respective Bayesian information criterion (BIC) as measure of how well each model captures the data, and converted the difference in BIC values into a Bayes factor for statistical interpretation (see section “Statistical Testing” for more details).

### Combined analysis of behavioral and EMREO data

We tested whether the state of the EMREOs at the time of Tone 2 affects how this tone is judged as follows: we relied on the same regression framework as above but included information about the EMREOs into the models. We did this in two ways, each testing a specific manner of how the EMREO may affect spatial hearing. One analysis probed the relevance of the direction of the EMREO deflections on systematic biases in the lateralization judgments; another analysis probed the influence of the magnitude of EMREO deflections on the accuracy of these judgments. The first analysis assumes that the differential in- and outwards deflections of the two eardrums at any given moment (i.e. positive and negative deflections of the EMREO) may result in a systematic error of how spatial acoustic cues are transduced (c.f. Fig 4). Since the EMREOs in the two eardrums are not directly in anti-phase, the relative deflection of these two eardrums may induce a spatial bias that varies along the EMREO time course. This spatial bias should manifest as generally more left (or right) responses for one particular sign of the EMREO compared to the other sign, regardless of the sound direction (illustrated in Figure 4A). The second analysis assumes that the overall deflection of the two eardrums, being small at some times and large at others, may affect how faithfully spatial information is transduced. This may affect the overall accuracy of spatial lateralization judgements. This should manifest as more (accurate) left/right responses for left/right tones for small EMREO amplitudes compared to large EMREO amplitudes (Figure 4C).

As above, we implemented these analyses using regression models that either predicted trial-wise response accuracy or the spatial judgement based on sound direction, saccade direction and information about the EMREOs. Since the analysis of behavior regardless of EMREO revealed that tone latency did not affect how sounds are judged, we did not include latency as a factor in these analyses. The EMREO values were determined as follows: for each participant we derived the trial-averaged EMREO separately for each ear and saccade direction. To determine the state of the two EMREOs for a given trial, we extracted the latency of Tone 2 for this trial, selected the EMREOs for the respective saccade direction, and used the continuous values of the left and right ear EMREOs at this latency (or the absolute value of this, depending on the analysis). To ensure a meaningful definition of the EMREO we focused these analyses only on trials in which the second tone was presented between 0 ms and 80 ms from saccade onset, i.e., the period containing the prominent EMREO signal. Given the pseudorandom presentation time of the tones relative to saccade onset, the resulting number of available trials differed between participants. The final dataset comprised data from 26 participants with 445.6 +/-12.4 trials (mean +/-SEM) in the free-field task and 21 participants with 346.7 +/-35.7 trials each in the in-ear task.

To directly demonstrate the participant-wise influence of EMREO sign and amplitude in line with this regression analysis, we included the expected effects and the actual data in Figure 4. Panel A illustrates the nature of the effect expected for the EMREO sign, and Panel B shows the difference in choice lateralization between the two EMREO signs. To derive this, we divided the trials for each participant based on the left-ear EMREO sign and computed a measure of overall choice directional bias for trials of each EMREO sign: this bias was obtained after normalizing the single-trial choice so that -1 corresponds to left and +1 to right. We then averaged this index across trials, with a value of 0 reflecting no bias and values trending to -1/+1 a bias towards left/right choices. Similarly, Panels C and D show the expected and actual effects for an influence of EMREO amplitude. Panel D shows a measure of the directional strength of participants choices for small and larger left-ear EMREO amplitudes (here implemented by a median split across all trials for each participant). This was obtained after grouping trials by the EMREO amplitude and computing the difference between the average normalized right minus left responses. A directional strength of 2 would correspond to perfect sound lateralization, and strength of 0 to no difference between tone directions. Splitting trials by the respective right ear EMREOS resulted in comparable figures.

### Statistical Testing

The statistical tests concerned the predictive power added by particular factors in the regression model (over their absence) and the significance of the slopes of individual predictors themselves. The added predictive power was measured using the difference in BIC values, which we converted to Bayes factors (Wagenmakers, 2007; Kass and Raftery, 1995). These Bayes factors (BF) then reflect the evidence in favor or against that a specific variable is predictive of the behavioral metric under consideration. In addition to reporting the individual BFs, we evaluate these qualitatively using the metric suggested by Raftery (1995): specifically, we consider BF below 3 as weak evidence, BF between 3 and 20 as positive, BF between 20-150 as strong, and BF above 150 as very strong evidence. Bayes factors larger than 1 indicate evidence that a given predictor adds predictive power, while values smaller than 1 indicate evidence against this hypothesis (Wagenmakers, 2007).

## Results

### Auditory Lateralization Performance Depends on the Congruency of Saccade Direction and Sound Direction

We first analyzed participants’ lateralization performance independent of the EMREOs. Using regression models, we probed how the sound direction, the saccade direction and the relative timing of the tones to saccade onset affected participants’ behavior. These models were fit to the trial-wise data and were used to predict response times, response accuracy and the actual left/right choices. From these models we obtained an estimate of whether a given predictor (e.g. saccade direction) adds significant predictive power to the model (derived in form of a Bayes factor, see Materials and Methods), and we investigated the significance of the individual predictors themselves.

For the free-field task (n=26) this revealed that the left/right choices were shaped by both the sound direction and the saccade direction (combined BF>10^5^; very strong evidence for an effect) but not the tone latency (BF=0.046; strong evidence against an effect). Both sound direction and saccade direction were significant predictors in the model (Table 1). Hence, participants’ judgments of sound direction were shaped not only by the acoustic signal but also by the saccade direction, in line with previous studies on how saccades shape auditory spatial perception. In line with this, response accuracy was shaped by the interaction of sound direction and saccade direction (BF>10^5^; very strong evidence for an effect) but not by tone latency (BF=0.048; strong evidence against an effect; see Table 1). Finally, response times revealed a significant influence of latency (BF=35.7; strong evidence for an effect) but not of saccade direction or sound direction (combined BF=0.0005; very strong evidence against an effect; Table 1). For the in-ear task (n=21) we obtained qualitatively similar results. Lateralization responses were shaped by both the sound and saccade directions (combined BF>10^5^; very strong evidence for an effect) but not latency (BF=0.058; moderate evidence against an effect; Table 2). Response accuracy was again shaped by the interaction of sound direction and saccade direction (combined BF>10^5^; very strong evidence for an effect) but not latency (BF=0.030; strong evidence against an effect; see Table 2). Finally, response times revealed no clear influence of latency (BF=0.59) and no influence of saccade direction or sound direction (combined BF =0.0013; very strong evidence against an effect; Table 2).

**Table 1.** Regression results for the behavioral data - Free-field task (n=26). The table reports the slopes (beta), their standard deviation (SD), and associated p-values. SoundDir:SaccDir denotes the interaction between sound and saccade directions. * represents p < 0.01

|  |  | beta | SE | p |
| --- | --- | --- | --- | --- |
| **Choice** | Intercept | 0.20219 | 0.13687 | 0.13987 |
|  | Sound Direction | 0.20724 | 0.066901 | 0.0019957* |
|  | Saccade Direction | 1.0543 | 0.067465 | 3.0176e-50* |
|  | SoundDir:SaccDir | 0.0074279 | 0.066906 | 0.91162 |
| **Accuracy** | Intercept | 0.539 | 0.012615 | 4.2311e-243 |
|  | Sound Direction | 0.044135 | 0.012615 | 0.00048471* |
|  | Saccade Direction | -0.0010482 | 0.012615 | 0.93379 |
|  | SoundDir:SaccDir | 0.23363 | 0.012615 | 1.5461e-67* |
| **Response Time** | Intercept | 0.95412 | 0.027298 | 4.843e-185* |
|  | Sound Direction | -0.0022666 | 0.0037313 | 0.54367 |
|  | Saccade Direction | -0.0084124 | 0.0037447 | 0.024857 |
|  | Latency | -0.00057003 | 0.00015058 | 0.00016103* |
|  | SoundDir:SaccDir | 0.0018502 | 0.003732 | 0.62015 |

**Table 2.** Regression results for the behavioral data - In-ear task (n=21). The table reports the slopes (beta), their standard error (SE), and associated p-values. SoundDir:SaccDir denotes the interaction between sound and saccade directions. * represents p < 0.01

|  |  | beta | SE | p |
| --- | --- | --- | --- | --- |
| **Choice** | Intercept | -0.046319 | 0.21047 | 0.82585 |
|  | Sound Direction | 0.64874 | 0.070481 | 1.5339e-19* |
|  | Saccade Direction | 0.99283 | 0.071527 | 1.1444e-40* |
|  | SoundDir:SaccDir | 0.057752 | 0.070004 | 0.40955 |
| **Accuracy** | Intercept | 0.63667 | 0.041644 | 3.0166e-48 |
|  | Sound Direction | -0.01099 | 0.012679 | 0.38624 |
|  | Saccade Direction | 0.0062633 | 0.012735 | 0.62295 |
|  | SoundDir:SaccDir | 0.20306 | 0.012678 | 2.3274e-52* |
| **Response Time** | Intercept | 0.94489 | 0.035573 | 4.9085e-122* |
|  | Sound Direction | 0.00021544 | 0.0038135 | 0.95496 |
|  | Saccade Direction | 0.0074125 | 0.0038419 | 0.053923 |
|  | Latency | -0.00048097 | 0.00019465 | 0.013616 |
|  | SoundDir:SaccDir | -0.006827 | 0.0038093 | 0.073356 |

While the group-level accuracy was above chance in both tasks (as indicated by the significant Intercepts for accuracy in both tasks; Tables 1 and 2) the overall accuracy was higher in the in-ear task (free-field: 0.55 +/-0.022; in-ear: 0.62 +/-0.021; two-sample t-test t=2.2, p=0.031).

These data are illustrated in Figure 2, which shows the participant-wise response accuracy and response times. To illustrate the influence of saccade direction, the data is shown for all trials, and for trials split by the congruency of saccade direction and sound direction. This directly illustrates that accuracy was higher for trials in which saccades direction and sound direction were congruent. The data is also shown as a function of the latency of Tone 2 relative to the saccade onset (trials were grouped in 20 ms time bins, 0ms corresponds to saccade onset). Trials in which Tone 2 was presented slightly before the saccade onset (time bin at -20 ms) were included as well. This reveals largely comparable performance for tones presented slightly prior to and during a saccade.

**Figure 2.**
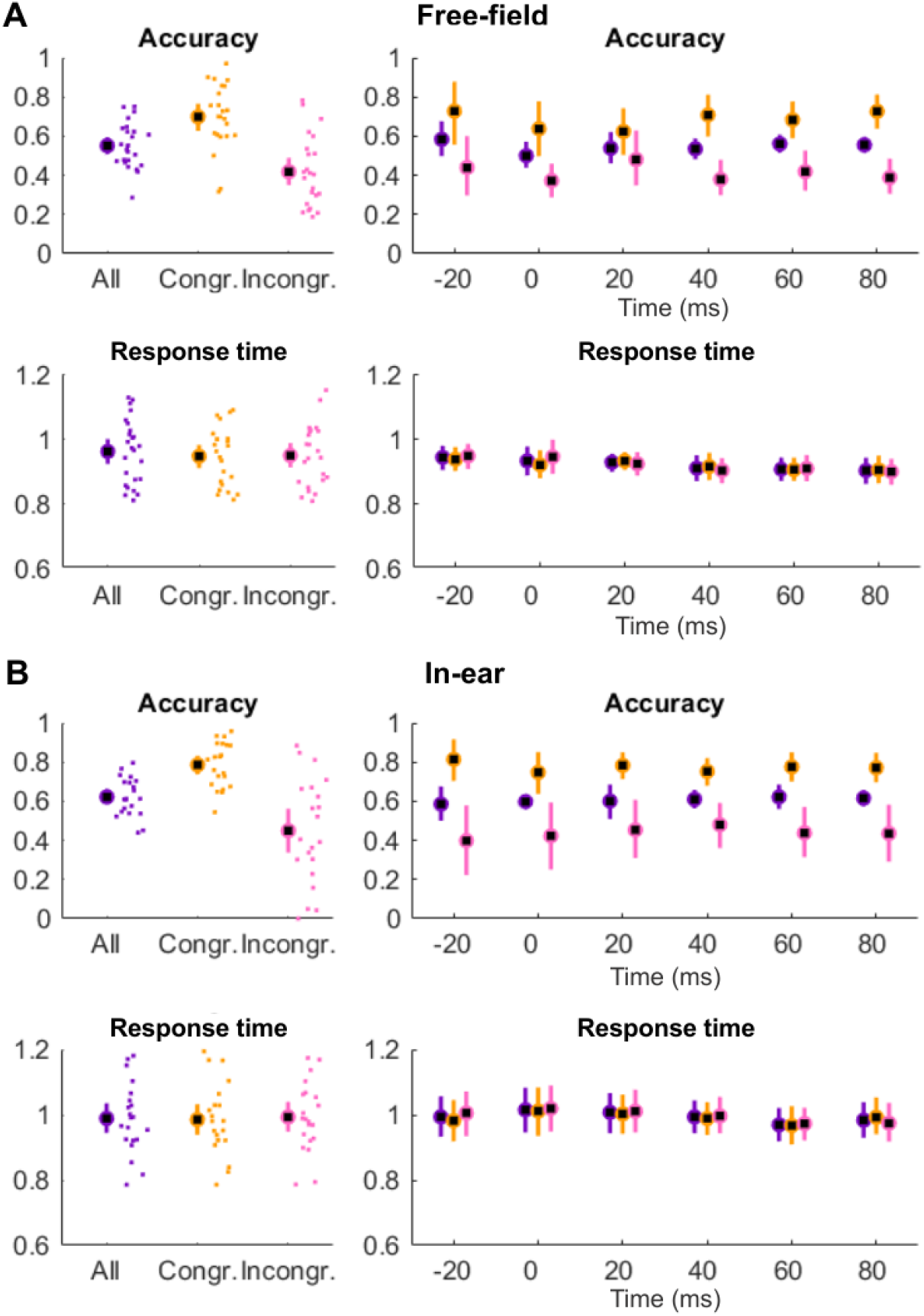
Behavioral data regardless of EMREOs. The left panels display participant-wise accuracy and response times for the sound direction for all trials (purple), and trials split by congruency between sound direction and saccade direction (congruent trials, yellow; incongruent trials, pink). Response times are in seconds. The right panels display the data as a function of the latency between Tone 2 and saccade onset (0 ms). Trials were grouped based on their respective tone latency using 20 ms bins (centered on the indicated time points). Since not all participants had sufficiently many trials for each of the time bins used for display here, the effective n differs between bins. Panel (**A**) shows the results for the free-field task, panel (**B**) for the in-ear task. The error bars indicate group means and 95th percentile bootstrap confidence intervals. Small dots indicate individual participant data.

### Group-level EMREOs

Figure 3A shows example data from one individual illustrating the left-ear EMREO together with the times of the key events during individual trials: the jump of the fixation target (gray), the presentation of the first tone (Tone 1; central reference, green) and the second tone whose direction participants had to judged (Tone 2; probe, pink). Given the trial-to-trial variability of saccade latencies, our experiment design effectively resulted in different and variable latencies of Tone 2 relative to the saccade onset. The eye velocity traces are shown for the individual example (Figure 3A, below the EMREO trace) and for the group-average for both tasks in (Figure 3B), illustrating the relation between the EMREO time course and saccade dynamics.

**Figure 3.**
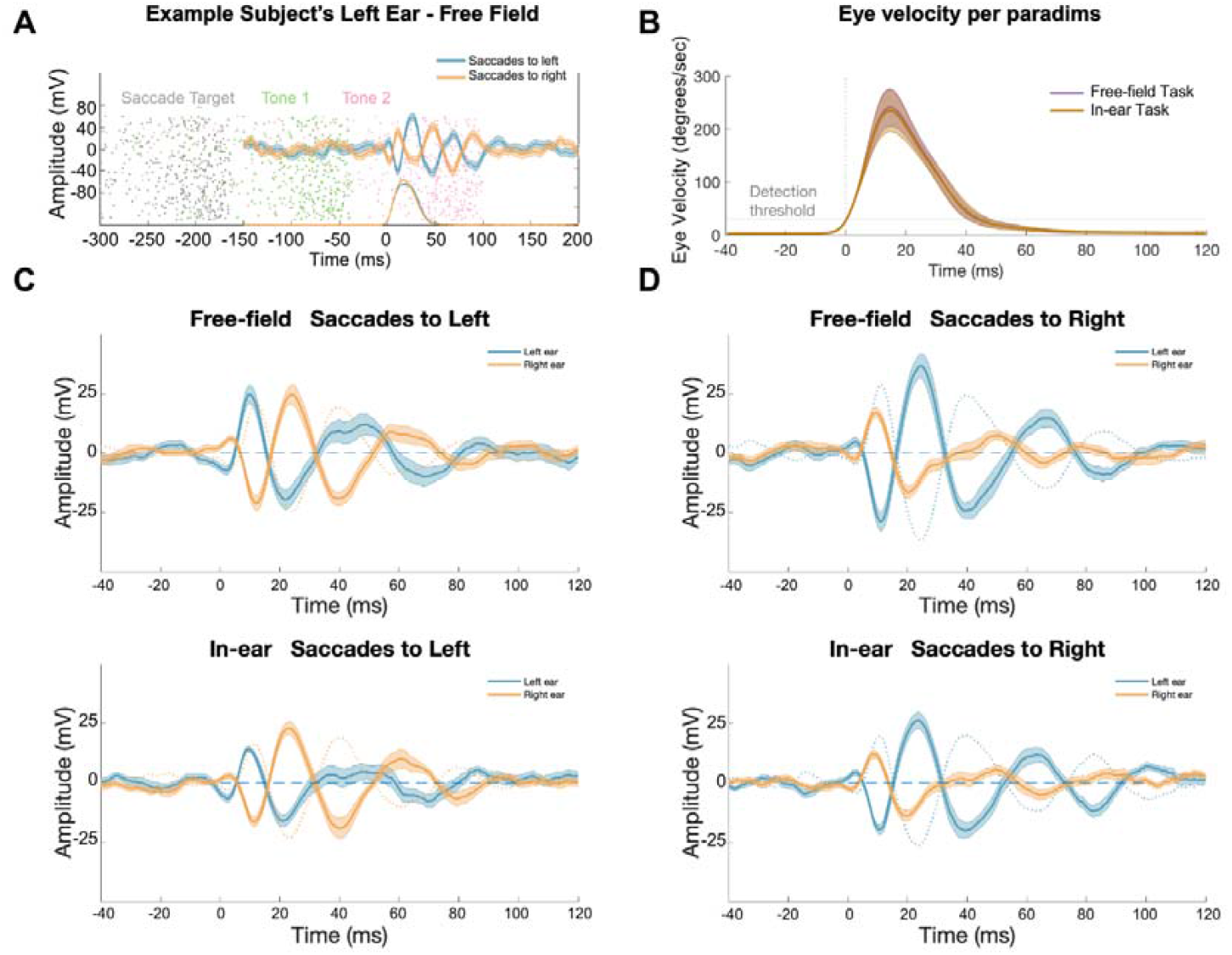
Example and Group-Level EMREOs. (**A**) shows example data for one participant, illustrating the EMREO in the left ear together with eye velocity during the free-field task. Dots display the presentation times for the saccade target (grey), the first tone (green) and the second tone (pink) in individual trials. (**B**) shows group-level eye velocity for the free-field (purple) and in-ear (brown) tasks, averaged across participants (mean and SEM across participants). (**C**) and (**D**) show group-average EMREOs for the free-field (n=26, 445.6 +/-12.4 trials) and in-ear (n=21, 346.7 +/-35.7 trials) tasks. EMREO time courses are shown for saccades to the left and right direction, split by ears (left ear traces in blue, right ear traces in orange). Thicker lines represent the mean EMREO, and shading reflects the SEM. For each panel, the contralateral EMREO with inverted sign is displayed as a dotted line for comparison with the corresponding ipsilateral trace. Time 0 ms is the detected saccade onset.

The EMREO signals obtained during the two auditory tasks followed the pattern reported in previous studies. Figure 3C and 3D show the group-level and trial-averaged EMREO time courses for both tasks. The time course of these reveals multiple deflections of alternating signs for each ear and saccades in the two directions, with prominent peaks between 10 and 60 ms following saccade onset. Importantly, within each ear the time course exhibits deflections of opposing signs for saccades in ipsi- and contralateral directions. However, for each ear the EMREO differs between ipsi- and contralateral saccades not only in sign but also the precise time course, as reported previously (Sotero Silva et al., 2025). As a result, for a given saccade the EMREOs in the two ears are not in strict anti-phase. Assuming that the EMREO time course directly reflects movements of the eardrums, this suggests that the relative movement of the left and right eardrums is not constant but changes over time following saccade onset. This leaves the possibility that spatial sound encoding is directly shaped by the combined state of the two eardrums in a manner that depends on the relative timing of a sound to the two EMREOs.

### Auditory Lateralization Performance Is Not Affected by Momentary EMREO Deflections

We performed two analyses to test whether the state of the tympanic membrane at the time of the second tone, as reflected by the EMREO signal, is relevant for participants’ performance. One analysis probed whether the direction of EMREO deflection predicts how participants judge the tone’s lateralization; thus, whether the direction of the ERMEO deflection induces response biases towards left or right responses. A second analysis probed whether the magnitude of the two EMREOs at the time of a tone predicts how accurately participants judge the tone’s lateralization. Each analysis then tests a specific manner in which movements of the eardrum reflected by the EMREO could affect spatial hearing.

Regarding the first analysis, the data suggest that the direction of the ERMEO deflection does not induce a specific spatial bias towards either left or right choices. For the free-field task we obtained very strong evidence against the added predictive power of the EMREO deflections on response choice (BF=0.0051) and neither EMREO was a significant predictor (Table 3). For the in-ear task the result was the same (BF=0.0043; Table 3).

**Table 3.** Regression results for the analysis testing and effect of EMREO sign on response choice. The table reports the slopes (beta), their standard error (SE), and associated p-values. Note that since the above results indicated that neither the interaction of sound and saccade direction nor the tone latency were significant predictors of choice, we did not include these as predictors in this analysis here. * represents p < 0.01

|  |  | beta | SE | p |
| --- | --- | --- | --- | --- |
| **Free-field Task** | Intercept | 0.18385 | 0.14459 | 0.2038 |
|  | Sound Direction | 0.25952 | 0.070998 | 0.00026928* |
|  | Saccade Direction | 1.0624 | 0.072365 | 1.2793e-44* |
|  | EMREO - left ear | 0.0071083 | 0.004034 | 0.078333 |
|  | EMREO - right ear | 0.0016261 | 0.0046341 | 0.72574 |
| **In-ear Task** | Intercept | -0.030568 | 0.20515 | 0.88158 |
|  | Sound Direction | 0.65864 | 0.074426 | 3.6008e-18* |
|  | Saccade Direction | 1.012 | 0.075772 | 1.0052e-37* |
|  | EMREO - left ear | 0.0039772 | 0.0048874 | 0.41596 |
|  | EMREO - right ear | 0.006348 | 0.0036941 | 0.086014 |

Regarding the second analysis, the data also do not suggest the magnitude of the EMREOs has an influence on how accurately participants judged the tone’s location. Neither for the free-field task nor for the in-ear task did the EMREO magnitude add predictive power (BF=0.0014 and BF=0.0012) nor was the EMREO a significant predictor (Table 4).

**Table 4.** Regression results for the analysis testing an effect of EMREO magnitude on response accuracy. The table reports the slopes (beta), their standard error (SE), and associated p-values. Note that since the above results indicated that tone latency was not a significant predictor of accuracy, we did not include this as a predictor in this analysis here.* represents p < 0.01

|  |  | beta | SE | p |
| --- | --- | --- | --- | --- |
| **Free-field Task** | Intercept | 0.15961 | 0.16467 | 0.33262 |
|  | Sound Direction | 0.22307 | 0.070799 | 0.0016737* |
|  | Saccade Direction | 0.027992 | 0.07569 | 0.71159 |
|  | EMREO - left ear magnitude | 0.00098227 | 0.0060585 | 0.87123 |
|  | EMREO - right ear magnitude | 0.0061996 | 0.0074418 | 0.40498 |
|  | SoundDir:SaccDir | 1.0511 | 0.071048 | 3.2416e-45* |
| **In-ear Task** | Intercept | 0.72848 | 0.27417 | 0.0080017 |
|  | Sound Direction | -0.054704 | 0.073573 | 0.45732 |
|  | Saccade Direction | 0.05254 | 0.077161 | 0.49607 |
|  | EMREO - left ear magnitude | -0.0047829 | 0.0072045 | 0.50692 |
|  | EMREO - right ear magnitude | 0.0014347 | 0.0048983 | 0.76966 |
|  | SoundDir:SaccDir | 0.99504 | 0.074374 | 7.7067e-38* |

To illustrate the participant-wise data underlying these results, Figure 4A,B shows the relation between EMREOs and the behavioral data for individual participants in the tasks. If a deflection of an EMREO would systematically bias participants to judge a sound towards one particular side, one would expect to see the following in the data (Figure 4A): the figure color-codes the direction of EMREO deflection and a systematic bias with EMREO direction should manifest in more responses to one side (e.g. left) for one EMREO sign (in the example yellow) and more responses to the other (here right) for the other sign (here purple color), regardless of the actual stimulus lateralization. However, such systematic patterns are not reflected in the actual data, in line with the null result from the statistical analysis (Figure 4B). Alternatively, if the presence of a strong EMREO deflection (large absolute EMREO) would deteriorate sound transmission this could be expected to result in lower response accuracy and hence a smaller difference of the average left /right judgement compared to smaller EMREO deflections. This behavior is illustrated in Figure 4C, illustrating more accurate responses for the purple condition (i.e. more left responses for left stimuli and more right responses for right stimuli) than for the yellow condition, where the numerical difference between choices for each stimulus side are smaller. Again, such an effect is not visible in the actual data (Figure 4D).

**Figure 4.**
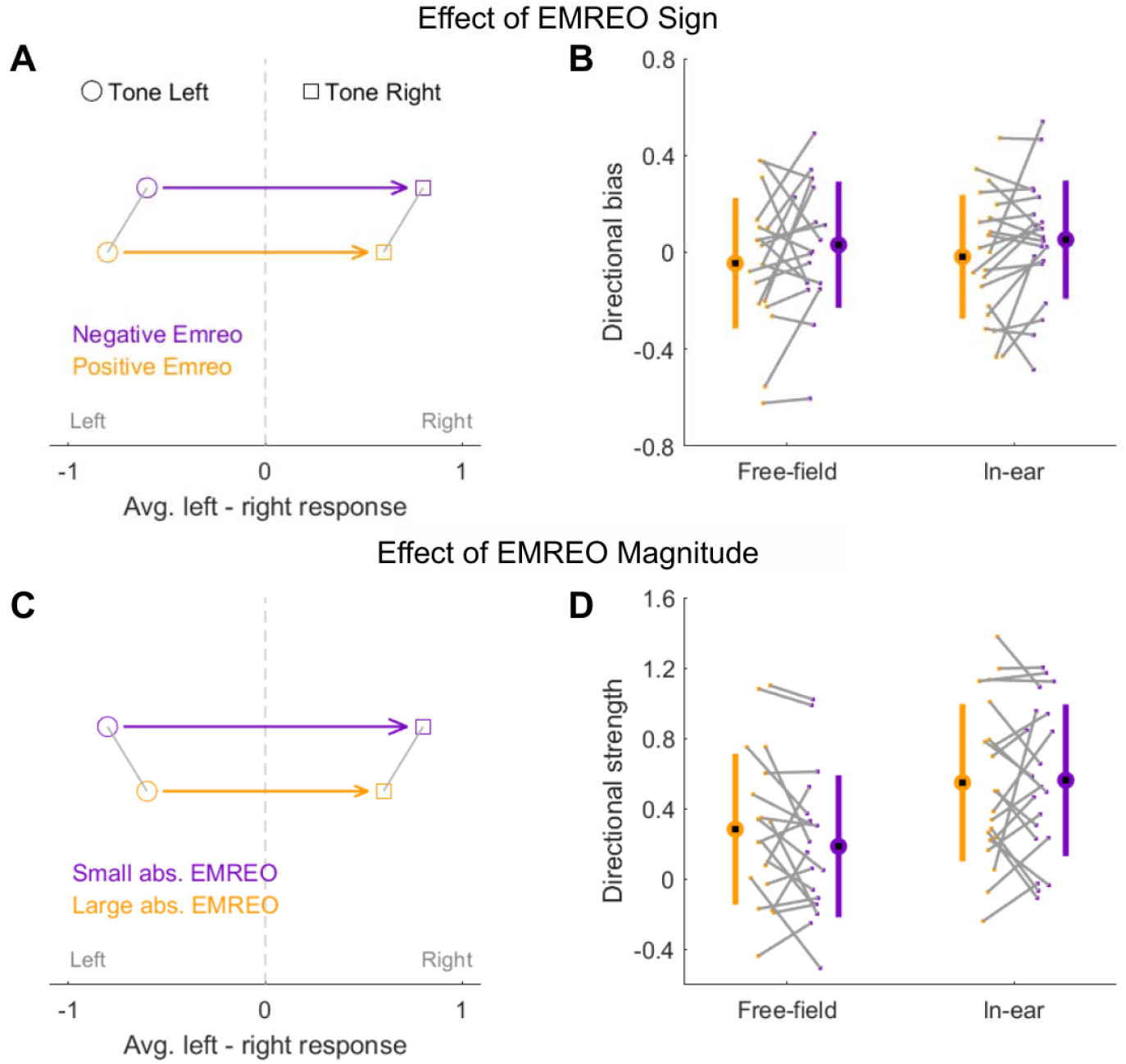
Hypothetical and actual effects of EMREO sign and magnitude on sound lateralization behavior. **(A**) shows a potential effect of EMREO sign on the fraction of left/right choices. The schematic illustrates choice behavior shown as a direction index with -1 corresponding to 100% left responses and +1 to 100% right responses for a given set of trials. If the sign of the EMREO would induce a direction bias, one could expect more right responses for one sign (here purple) compared to the other sign (here yellow) regardless of the lateralization of the tone (circle, square). This overall shift in response bias was quantified for each participant as a directional bias index and averaged across both tone lateralization. (B)shows the resulting distribution of directional biases for both EMREO signs and tasks. In line with the regression analyses, the distributions of directional biases overlap strongly. (**C**) shows a potential effect of EMREO magnitude (absolute value). One could expect that a small EMREO magnitude (purple) would distort the accuracy of lateralization choices less than a large EMREO magnitude (yellow). Hence, the difference in left/right choice behavior between left and right tones should be larger in the purple compared to the orange condition. (**D**) shows the effect in the actual data in the form of a directional strength index. This was computed as a difference in proportion of right minus left choices. As expected, and based on the regression results, the two distributions overlap.

## Discussion

Eye movement-related eardrum oscillations (EMREOs) describe pressure changes in the outer ear following saccadic eye movements. Despite several recent studies, their origin and function remain unclear (Sotero Silva et al., 2025; King et al, 2023; Lovich et al., 2023a, 2023b; Gruters et al., 2018). The generation of the EMREOs has been postulated to involve the two middle ear muscles (MEMs, the tensor tympani and stapedius) and possibly the outer hair cells (Lovich et al., 2023b; Gallagher, Diop and Olson, 2021; Gruters et al., 2018). The MEMs implement critical functions in protecting hearing by increasing the stiffness of the ossicular chain (Edmonson et al., 2022; Racca et al., 2022; Lawrence, 1965). Small changes in the stiffness have been shown in *ex-vivo* studies to alter the phase of the transmitted sound and thereby distort the encoding of temporal cues that are important for acoustic lateralization (Cho, Ravicz and Puria, 2023). Importantly, such changes in stiffness could also be induced during the EMREOs. Hence, a sound arriving at the eardrum in a stronger EMREO deflection, or a deflection towards one particular side, may be transduced differently compared to a sound received during a different portion of the EMREO (O’Connor, Cai and Puria, 2017). Consequently, one could expect an influence of the EMREO on encoding the location of acoustic signals. We tested this experimentally by probing whether the presentation of sounds at different phases of the EMREO affects how well their location can be judged. However, consistently across two tasks the evidence does not point in favor of such an effect.

We take this to suggest that either the interpretation of the EMREO signal as directly reflecting movements of the tympanic membrane is incorrect. Or, alternatively, that the effective changes at the eardrum are too short-lived or have only negligible perceptual effects. The assumption that the temporally extended counter-phase pattern of the EMREO reflects movements of the tympanic membrane on the same time scale is central to the logic of the present experiments. Under this assumption, and the assumption that MEMs are involved in the EMREO, it is reasonable to expect a difference in behavioral performance between sounds presented at different deflections of the EMREO. Still, the lack of behavioral consequences observed here is incompatible with these assumptions.

### Understanding the Origin of the EMREO Signal

The EMREO waveform is likely associated with the coordinated contraction of the tensor tympani and stapedius, with a possible contribution from the outer hair cells (Gruters et al., 2018; Lovich et al., 2023b). However, the EMREO signal is an acoustic signal recorded using in-ear microphones and only reflects pressure changes. Whether the tympanic membrane indeed moves directly in proportion to the EMREO signal remains still unknown. One possibility is that the tympanic membrane only moves very briefly, or changes its tension, and the resulting temporally extended signal recorded is only an acoustic reverberation resulting from this brief change. In this interpretation the EMREO could still be a peripheral signature of a central motor command involved in oculomotor function, but that command may translate into only a brief action on the tympanic membrane (Lovich et al. 2023b, Sotero Silva et al., 2025). This does not rule out that the EMREO has a detectable impact on spatial sound encoding and perception, or that it plays a role in combining multisensory spatial signals (Gruters et al, 2018; Murphy et al., 2020; Lovich et al., 2023a). Nevertheless, this action would then be limited to a short time directly around the actual action of the MEMs, e.g., directly at saccade onset, and would hence only affect very brief sounds impinging on the ear at precisely this short-lived time point. Whether such a short-lived effect exists remains unclear based on the present data and would require an experimental paradigm that precisely aligns sounds to saccade onsets at the millisecond level.

Another possibility is that the eardrum moves in line with the EMREO signal, but the actual changes in membrane stiffness and sound transduction delays caused by the MEMs are irrelevant for sound lateralization. Relative eardrum stiffness, modulated, for example, by the acoustic reflex, alters the perceived properties of sounds (Racca□et□al.,□2022;□ Avan□et□al.,□2000), such as attenuating low-frequency transmission and shortening the middle-ear transmission group delay (O’Connor, Cai and Puria,□2017). At the same time, the magnitude of an individual’s EMREO may be related to eardrum compliance, whereby lower compliance, therefore larger stiffness, is associated with a higher EMREO peak amplitude (Sotero□Silva et al.,□2025). Whether the action of the MEMs during the EMREO has a meaningful impact on sound transduction under everyday settings still is unclear, as previous data supporting sound transduction being mediated by MEMs tension were obtained *ex-vivo* (Cho, Ravicz and Puria, 2023). However, obtaining direct information on the actual movement of the eardrum in humans *in vivo* during task performance is a challenge (Huber et al., 2026) and would require measurements that are not affected by concurrently presented acoustic stimuli (e.g., using laser doppler vibrometry).

Alternatively, the eardrum could move for a prolonged time in line with the EMREO signal, but this may impact sound lateralization performance only under conditions different than those tested here. For example, it could be that any relation of EMREOs to spatial hearing emerges only for very large or long-lasting saccades. While some studies suggest that the EMREO amplitude may scale with saccade size, the direct evidence visible in previous studies suggests that any such scaling is limited and may asymptote for saccades larger than about 10 to 15 degrees (Fig 2D and F. Gruters et al., 2018). While the precise scaling of the EMREO with saccade size remains to be studied in detail, the existing evidence casts doubts on the possibility that the EMREO induced by very large saccades (>20 degrees) has a more pronounced effect than an EMREO for saccades around 5 to 10 degrees, as tested here.

Further, it is possible that the movement of the eardrum reflected in the EMREO matters only for sounds other than those tested here. The present experiments relied on a single brief tone of 1200 Hz. The tone was presented in a free-field condition and in an in-ear condition, with the latter only varying the stimulus ITDs to generate the percept of different azimuthal positions. ITDs are mostly relevant for estimating azimuthal positions for low frequency sounds, i.e., below 2000 Hz (Azevedo et al., 2024; Wightman and Kistler, 1992). While the present sounds fall in this range, it could be that the perceptual effect becomes relevant only at much lower sound frequencies and future studies may consider a wider range of sounds.

Finally, the laboratory setting used to study EMREOs does not replicate the multifaceted conditions present in natural environments, where EMREOs may possibly play a more pronounced role for hearing. For example, the studies to date rely on a fixed head position and eye movements that do not bear any relation to the acoustic task performed at the same time. However, if the EMREO reflects a mechanism genuinely involved in combining visual and acoustic signals, one may expect this mechanism to operate in a task-specific manner. Many everyday conditions involve multiple visual and acoustic stimuli, and attention and task-relevance are critical factors shaping multisensory integration under such conditions (Noppeney, 2021). To address this, future studies could employ tasks such as tracking moving sounds, or tasks detecting subtle changes in a sound’s properties to increase ecological validity.

### Congruency of Saccade and Sound Direction Modulates Performance

While we did not find evidence that the state of the eardrum as reflected by the EMREO affects spatial sound lateralization judgements, our data show that the congruency between the saccade direction and the relative location of two tones affects how this relative location is judged. In the present paradigm the probe sound locations (tone 2) were either displaced in the same direction or the opposite of the saccade and participants accuracy was higher in congruent trials. In support of this, the regression analyses confirmed that both the sound direction and the saccade direction significantly influenced participants left/right choices.

This interaction of oculomotor behavior and spatial auditory perception is consistent with previous studies showing that both perceived sound locations and judgements of auditory motion are biased towards visual saccade targets (Krüger et al., 2016; Klingelhoefer and Bremmer, 2009; Pavani et al., 2008; Binda et al., 2007). One of these previous studies directly probed auditory judgements at different latencies relative to saccade onset, and found that the resulting perceptual biases are fairly stable relative to the time course of a saccade (Klingelhoefer and Bremmer, 2009). This result is also in line with our conclusion that the EMREO time course has no clear impact on spatial hearing. Together with studies showing that auditory space is perceived differently depending on where participants fixate their gaze (Lewald and Ehrenstein, 1996; Lewald, 1997; Getzmann, 2002; Razavi et al., 2007; Maddox et al., 2014) this points to a profound interaction between the oculomotor system, general attentional control and spatial perception. While recent studies on EMREOs uphold a potential role of the underlying neural processes in shaping cross-modal attention and or sensorimotor integration, their precise impact on spatial hearing and perception in general still remains to be fully understood.

## Author contributions

**Nancy Sotero Silva:** Performed experiments, Analyzed data, Interpreted results of experiments, Prepared figures, Drafted manuscript, Edited and revised the manuscript. **Felix Bröhl:** Conceived and designed research, Analyzed data, Interpreted results of experiments, Drafted manuscript, Edited and revised the manuscript, Approved final version of the manuscript. **Christoph Kayser:** Conceived and designed research, Analyzed data, Interpreted results of experiments, Drafted manuscript, Approved final version of the manuscript.

## References

1. Sotero Silva N, Kayser C, Bröhl F. Unraveling eye movement-related eardrum oscillations (EMREOs): how saccade direction and tympanometric measurements relate to their amplitude and time course. Hearing Research 461: 109276, 2025. 10.1016/j.heares.2025.109276

2. King CD, Lovich SN, Murphy DLK, Landrum R, Kaylie D, Shera CA, Groh JM. Individual similarities and differences in eye-movement-related eardrum oscillations (EMREOs). Hearing Research 440: 108899, 2023. 10.1016/j.heares.2023.108899

3. Lovich SN, King CD, Murphy DLK, Abbasi H, Bruns P, Shera CA, Groh JM. Conserved features of eye movement related eardrum oscillations (EMREOs) across humans and monkeys. Philosoph. transact. Royal Soc. London Series B, Biol. sci., 378: 20220340, 2023a. 10.1098/rstb.2022.0340

4. Gruters KG, Murphy DLK, Jenson CD, Smith DW, Shera CA, Groh JM. The eardrums move when the eyes move: A multisensory effect on the mechanics of hearing. PNAS 115(6): E1309–E1318, 2018. 10.1073/pnas.1717948115

5. Lovich SN, King CD, Murphy DLK, Landrum RE, Shera CA, Groh JM. Parametric information about eye movements is sent to the ears. PNAS 120(48): e2303562120, 2023b. 10.1073/pnas.2303562120

6. Murphy DLK, King CD, Lovich SN, Landrum RE, Shera CA, Groh JM. Evidence for a system in the auditory periphery that may contribute to linking sounds and images in space. bioRxiv 2020.07.19.210864, 2020. 10.1101/2020.07.19.210864

7. Bröhl F, Kayser C. Detection of Spatially Localized Sounds Is Robust to Saccades and Concurrent Eye Movement-Related Eardrum Oscillations (EMREOs). Journal of Neuroscience 43(45): 7668–7677, 2023. 10.1523/JNEUROSCI.0818-23.2023

8. Culling JF, Akeroyd MA. F. Spatial hearing. In: The Oxford handbook of auditory science: hearing (online edition), edited by Plack CJ. Oxford Library of Psychology, 2010. 10.1093/oxfordhb/9780199233557.013.0006

9. Cho NH, Ravicz ME, Puria S. Human middle-ear muscle pulls change tympanic-membrane shape and low-frequency middle-ear transmission magnitudes and delays. Hearing Research 430: 108721, 2023. 10.1016/j.heares.2023.108721

10. Tasko SM, Deiters KK, Flamme GA, Smith MV, Murphy WJ, Jones HG, Greene NT, Ahroon WA. Effects of unilateral eye closure on middle ear muscle contractions. Hearing Research 424: 108594, 2022. 10.1016/j.heares.2022.108594

11. Harris LR, Lieberman L. Auditory stimulus detection is not suppressed during saccadic eye movements. Perception 25: 999–1004, 1996. 10.1068/p250999

12. Paire A, Vergilino-Perez D, Paeye C. Vertical shifts of visuospatial attention, not (eye) movements, affect auditory pitch discrimination. Journal of Vision 25(12): 6, 2025. 10.1167/jov.25.12.6

13. Lewald J, Ehrenstein WH. The effect of eye position on auditory lateralization. Experimental Brain Research 108: 473–485, 1996. 10.1007/BF00227270

14. Lewald J. Eye-position effects in directional hearing. Behavioural Brain Research 87(1): 35–48, 1997. 10.1016/S0166-4328(96)02254-1

15. Getzmann S. The effect of eye position and background noise on vertical sound localization. Hearing Research 169: 130–139, 2002. 10.1016/S0378-5955(02)00387-8

16. Klingenhoefer S, Bremmer F. Perisaccadic localization of auditory stimuli. Experimental Brain Research 198: 411–423, 2009. 10.1007/s00221-009-1869-3

17. Van Grootel TJ, Van Opstal AJ. Human Sound-Localization Behavior Accounts for Ocular Drift. Journal of Neurophysiology 103(4): 1927–1936, 2010. 10.1152/jn.00958.2009

18. Razavi B, O’Neill WE, Paige GD. Auditory spatial perception dynamically realigns with changing eye position. Journal of Neuroscience 27(38): 10249–10258, 2007. 10.1523/JNEUROSCI.0938-07.2007

19. Pavani F, Husain M, Driver J. Eye-movements intervening between two successive sounds disrupt comparisons of auditory location. Experimental Brain Research 189(4): 435–449, 2008. 10.1007/s00221-008-1440-7

20. Binda P, Bruno A, Burr DC, Morrone MC. Fusion of visual and auditory stimuli during saccades: a Bayesian explanation for perisaccadic distortions. The Journal of Neuroscience 27(32): 8525–8532, 2007. 10.1523/JNEUROSCI.0737-07.2007

21. Krüger HM, Collins T, Englitz B, Cavanagh P. Saccades create similar mislocalizations in visual and auditory space. Journal of Neurophysiology 115: 2237–2245, 2016. 10.1152/jn.00853.2014

22. World Health Organization. World report on hearing (Online). https://www.who.int/publications/i/item/9789240020481 [2021 March 3rd]

23. Brainard DH. The Psychophysics Toolbox. Spatial Vision 10(4): 433–436, 1997. 10.1163/156856897X00357

24. The MathWorks Inc. MATLAB version: 9.13.0 (R2022b), 2022. https://www.mathworks.com

25. SR Research Ltd. EyeLink 1000 Plus. [Apparatus and software], 2015. https://www.sr-research.com/eyelink-1000-plus/

26. American Speech-Language-Hearing Association (Online). (Central) auditory processing disorder—the role of the audiologist - Position Statement. www.asha.org/policy [2005]

27. Wagenmakers EJ. A practical solution to the pervasive problems of p values. Psychonomic Bulletin & Review 14(5): 779e804, 2007. 10.3758/BF03194105

28. Kass RE, Raftery AE. Bayes Factors. Journal of the American Statistical Association 90 (430): 773–795, 1995. 10.2307/2291091

29. Raftery AE. Bayesian model selection in social research. Sociological methodology 25: 111–196, 1995. 10.2307/271063

30. Gallagher L, Diop M, Olson, ES. Time-domain and frequency-domain effects of tensor tympani contraction on middle ear sound transmission in gerbil. Hearing Research 405: 108231, 2021. 10.1016/j.heares.2021.108231

31. Edmonson A, Iwanaga J, Olewnik Ł, Dumont AS, Tubbs RS. The function of the tensor tympani muscle: a comprehensive review of the literature. Anatomy & cell biology 55(2): 113–117, 2022. 10.5115/acb.21.032

32. Racca JM, Delgado RE, Gifford RH, Ramachandran R, Hood LJ. The Effects of Middle-ear Stiffness on the Auditory Brainstem Neural Encoding of Phase. Journal of the Association for Research in Otolaryngology 23(6): 859–873, 2022. 10.1007/s10162-022-00872-0

33. Lawrence M. Middle ear muscle influence on binaural hearing. Archives of Otolaryngology 82(5): 478–482, 1965.

34. O’Connor KN, Cai H, Puria S. The effects of varying tympanic-membrane material properties on human middle-ear sound transmission in a three-dimensional finite-element model. The Journal of the Acoustical Society of America 142(5): 2836–2853, 2017. 10.1121/1.5008741

35. Avan P, Büki B, Maat B, Dordain M, Wit HP. Middle ear influence on otoacoustic emissions. I: noninvasive investigation of the human transmission apparatus and comparison with model results. Hearing Research 140(1–2): 189–201, 2000. 10.1016/S0378-5955(99)00201-4

36. Huber A, Baselt B, Dobrev I, Prochazka L, Pfiffner F, Etter D, Peter-Siegrist N, Röösli C, Sim JH, Schär M. Methods matter: Current and future practices for middle ear mechanics laboratories. Hearing Research 470: 109505, 2026. 10.1016/j.heares.2025.109505

37. Azevedo ACPFO, Pellegrino TG, Peña JL, Marin B, Pavão R. Sufficiency of ITD for (biased) human auditory azimuthal perception. bioRxiv 2024.11.05.622135, 2024. 10.1101/2024.11.05.622135

38. Wightman FL, Kistler DJ. The dominant role of low-frequency interaural time differences in sound localization. The Journal of the Acoustical Society of America. 91: 1648–1661, 1992. 10.1121/1.402445

39. Noppeney U. Perceptual inference, learning, and attention in a multisensory world. Annual review of Neuroscience 44(1): 449–473, 2021. 10.1146/annurev-neuro-100120-085519

40. Maddox RK, Pospisil DA, Stecker GC, Lee AK. Directing eye gaze enhances auditory spatial cue discrimination. Current Biology 24(7): 748–752, 2014. 10.1016/j.cub.2014.02.021

